# Biosynthesis of H-GDGTs linked to ocean oxygen deficiency

**DOI:** 10.1101/2023.10.20.562873

**Authors:** Yanan Li, Ting Yu, Xi Feng, Bo Zhao, Huahui Chen, Gregory T. Connock, Xiao-Lei Liu, Huan Yang, Jérôme Kaiser, Anna K. Wittenborn, Liang Dong, Fengping Wang, Hayden R. Anderson, Noah Z. Burns, Fuxing Zeng, Lizhi Tao, Zhirui Zeng

**Author notes:** **Corresponding authors:** Fuxing Zeng, Lizhi Tao, and Zhirui Zeng.

## Abstract

Archaeal membrane lipids GDGTs (glycerol dialkyl glycerol tetraethers) are biomarkers used for tracking Earth’s historical environmental changes. Among these GDGTs, the H-shaped GDGTs (H-GDGTs, or GMGTs) represent a less-explored and often overlooked subset, with its biosynthetic pathway and geological significance remaining elusive. Here, we identified the gene responsible for biosynthesizing H-GDGTs, which encodes to a radical *S*-adenosyl-L-methionine (SAM) enzyme, named as H-GDGTs bridge synthase (Hbs). Heterologous expression of the gene *hbs* in a methanogen, as well as *in vitro* activity assay using the purified Hbs enzyme were performed. Additionally, we found that the genes encoding Hbs are exclusively present in obligate anaerobic archaea genomes and the metagenomes obtained from oxygen-deficient environments, but not in oxic settings. The H-GDGTs lipids were also consistently enriched in the modern oxygen-deficient environments, and remarkably accumulated in ancient sediments during oceanic anoxic event-2 (OAE-2, ∼94 million years ago) period. Our findings indicate H-GDGTs holds significant promise as a novel biomarker for studying historical ocean oxygen deficiency supported by a well-established biological basis.

## Introduction

The depletion of oxygen in the ocean poses a significant threat to the diversity and productivity of marine ecosystems (*1*). Throughout Earth’s geological history, episodes of ocean anoxia, known as Ocean Anoxic Events (OAEs), have occurred repeatedly (*2*). These anoxic events have been recorded by distinctive geochemical signatures, which encompass total organic carbon (TOC), stable isotopes of essential elements (e.g., δ^13^C and δ^34^S), redox-sensitive trace metals (e.g., Mo, V, U), and specific lipid biomarkers (*3*).

Archaeal spanning membrane lipids GDGTs (Glycerol Dialkyl Glycerol Tetraethers), play a crucial role as lipid biomarkers in the reconstruction of paleoclimate changes, including OAEs (*4*). Recent advancements in unraveling the GDGTs biosynthetic pathway have laid a strong biological foundation for the effective utilization of GDGTs proxies. The key enzymes responsible for GDGTs biosynthesis, particularly the processes of C(sp^3^)-C(sp^3^) cross-linkage reaction (*5, 6*) and cyclization (*7*), belong to the radical *S*-adenosyl-L-methionine (SAM) superfamily. These enzymes utilize a radical SAM[4Fe-4S] cluster in combination with SAM to generate a super reactive 5’-deoxyadenosine radical (5’-dAdo•) via the reductive cleavage of SAM. The 5’-dAdo• then initiates diverse radical chemistries through either hydrogen atom (H-atom) abstraction or substrate adenosylation (*8*). In the biosynthesis of GDGTs, the orchestrated utilization of such high-energy radical proves highly advantageous for forming the challenging C(sp^3^)-C(sp^3^) linkage.

H-GDGTs (or GMGTs, Glycerol Monoalkyl Glycerol Tetraethers) constitute a subset of GDGT derivatives distinguished by an extra cross-linkage between two isoprenoid chains (*9*). These unique lipids have been identified in multiple thermophilic archaea and are thought to contribute to increased membrane rigidity (*10*). Nevertheless, the lack of knowledge regarding its biosynthesis, synthetic mechanisms, and comprehensive biological sources obscures its potential significance as biomarker (*11*).

## Results

### Discovery and identification of H-GDGTs bridge synthase *in vivo* and *in vitro*

The biosynthesis of H-GDGT is likely through generating C-C bridge of two isoprenoid chains on GDGT, likely a covalent bond between two methyl groups (*12*). Inspired by the radical SAM enzymes, including Tes (*5*) or its homologue MJ0619 (*6*) and GrsA/B (*7*), which can activate the completely inert sp^3^-hybridized carbon centers to accomplish a challenging C(sp^3^)-C(sp^3^) bond formation reaction, we searched the gene by employing the radical SAM as candidates. We compared radical SAM enzymes (pfam04055) between two archaeal groups: one group involving *Pyrococcus furiosus* and *Thermococcus guaymasensis* (*10, 13*) produces H-GDGTs, while the other group including *Pyrococcus yayanosii* and *Thermococcus kodakarensis* (*14*) does not. We successfully found a promising candidate (WP_062369926.1) from *T. guaymasensis*. This gene contains a canonical CX_3_CX_2_C motif which provides three cysteine (Cys) residues to ligate a RS [4Fe-4S] cluster for initiating radical SAM chemistry (Fig. S1). Therefore, it is most likely that this gene is responsible for forming H-GDGTs from GDGTs.

To test our hypothesis, we conducted *in vivo* studies using an engineered methanogen *Methanococcus maripaludis* as the host for heterologous expression. This particular methanogen harbors the *tes* gene (*5*) and can produce GDGT-0 as the substrate for H-GDGTs bridge synthase (Fig. 1A). Also we cloned and expressed the gene homolog of WP_062369926.1 from the methanogen *Methanothermococcus okinawensis* (*15*), known as METOK_RS04425, to improve the expression efficiency of exogenous protein in the methanogen host (Fig. 1B left). The clones were cultured at 37°C for 7 days and then harvested for lipid analysis with reversed-phase ultra-high performance liquid chromatography mass spectrometry (UPLC-MS/MS). The results clearly showed the production of a lipid compound with *m/z* 1300; and due to the covalent bond between the two isoprenoid chains, the Tandem MS/MS spectra only show H-GDGT-0 with [M+H]^+^ ions and the lost water and glycerol (Fig .1C). This analysis confirms the *in vivo* formation of H-GDGT-0, indicating that the radical SAM enzyme encoded by METOK_RS04425 is responsible for H-GDGTs biosynthesis (Fig. 1A). Therefore, we name this enzyme as H-GDGTs bridge synthase (Hbs).

**Fig. 1.**
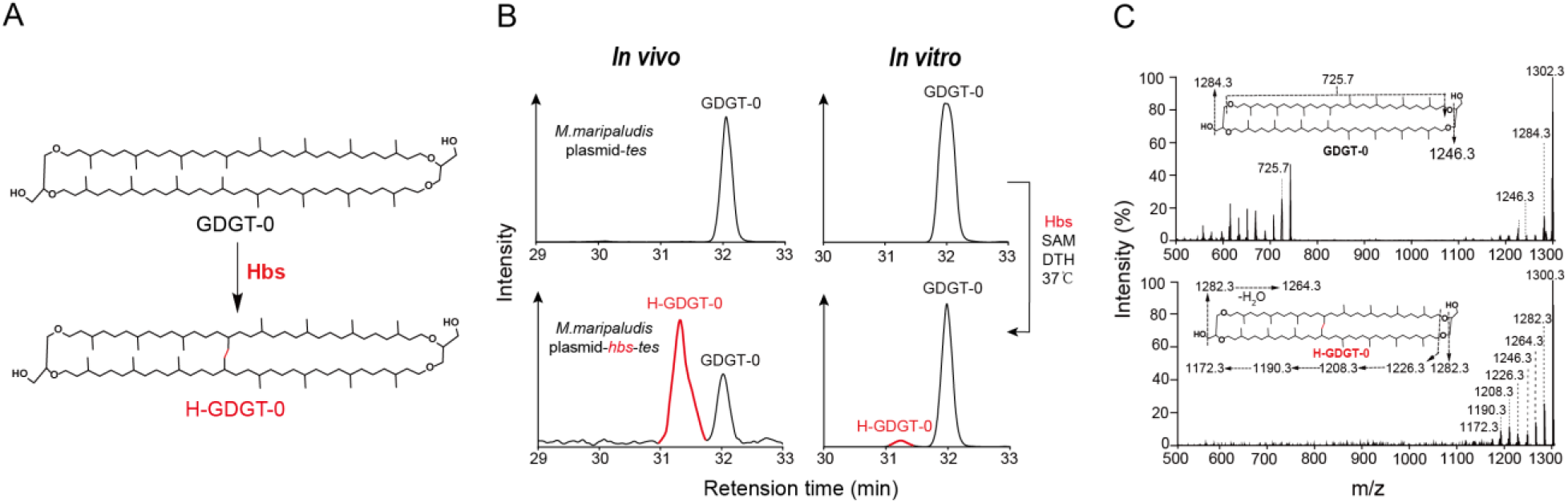
Identification of H-GDGTs bridge synthase *in vivo* and *in vitro* assays. (A) H-GDGT-0 biosynthesis pathway. (B) UPLC-MS extracted ion chromatograms of lipids extracts from *M. maripaludis* with plasmid pMEV4-*tes* showing production of GDGT-0, and with plasmid pMEV4-*hbs*-*tes* producing H-GDGT-0; and *in vitro* biochemical reaction showing to synthesize H-GDGT-0 with the presence of purified Hbs protein, GDGT-0, SAM and DTH at 37°C for 72 hours. (C) MS/MS spectra of GDGT-0 and H-GDGT-0.

To confirm the function of Hbs, we purified Hbs enzyme and performed *in vitro* studies. Recombinant Hbs from *M. okinawensis* was heterologously expressed in *Escherichia coli* BL21(DE3) and purified anaerobically using *Strep*-Tactin affinity chromatography. A mutant strain of *Sulfolobus acidocaldarius*, which produces GDGT-0 as its dominant membrane lipid (*7*), was employed for isolating the substrate GDGT-0 using preparative LC-MS. Then an *in vitro* activity assay was performed by incubating Hbs with 30 equiv. of reductant dithionite (DTH), 30 equiv. of SAM and 10 equiv. of GDGT-0 at 37 °C anaerobically for 72 hours. The product of H-GDGT-0 was observed and confirmed, with a yield of approximately 4.8% (Fig. 1B right and Fig. S2). These results unequivocally demonstrate that Hbs catalyzes a C-C linkage between two isoprenoid chains on GDGT-0 to form H-GDGT-0.

To further elucidate the details of H-GDGT-0 biosynthesis, we employed electron paramagnetic resonance (EPR) spectroscopy to characterize the FeS cluster(s) in the purified Hbs. The EPR spectrum of the as-purified Hbs reduced by DTH shows a complex FeS signal (Fig. 2). Subsequent addition of SAM results in an immediate spectral change, suggesting SAM binds to the radical SAM [4Fe-4S]^+^ cluster and affects the electronic structure of cluster significantly. Additionally, we observed more than one FeS cluster signal in the spectra. In order to deconvolute the signal, we employed AlphaFold2 (*16*) to predict the structure of Hbs. The model showed that Hbs contains four domains, including an N-terminal domain comprised of six helices, a conserved radical SAM domain, a bridging region with two helices and a β-harpin motif, as well as a C-terminal SPASM domain (Fig. 2A and Fig. S3). By aligning with the atomic structure of SPASM-domain containing radical SAM enzyme SuiB (PDB: 5V1T) (*17*), three [4Fe-4S] clusters were demonstrated in our Hbs model, consistent with the complex FeS cluster signal observed in EPR spectra (Fig. 2B and Fig. S3). With the aid of the structural model, we prepared corresponding Hbs mutants by knocking out two [4Fe-4S] clusters, leaving only one cluster in each (Fig. 2E, F). The **g** tensors for the three clusters were clearly determined via spectral simulation, i.e. [2.051, 1.884, 1.817] for SAM-bound radical SAM cluster, [2.045, 1.930, 1.885] for AuxI cluster and [2.036, 1.924, 1.871] for the AuxII cluster (Fig. 2F). The g tensors are typical for both radical SAM cluster and the auxiliary [4Fe-4S] clusters in the SPASM-domain containing radical SAM enzymes (*18*). In addition, we did not observe a SAM response for both AuxI and AuxII clusters (Fig. S3), suggesting that SAM does not bind to the auxiliary clusters as it does for radical SAM cluster. This further corroborates the structural model showing the two auxiliary clusters are fully ligated by four Cys residues.

**Fig. 2.**
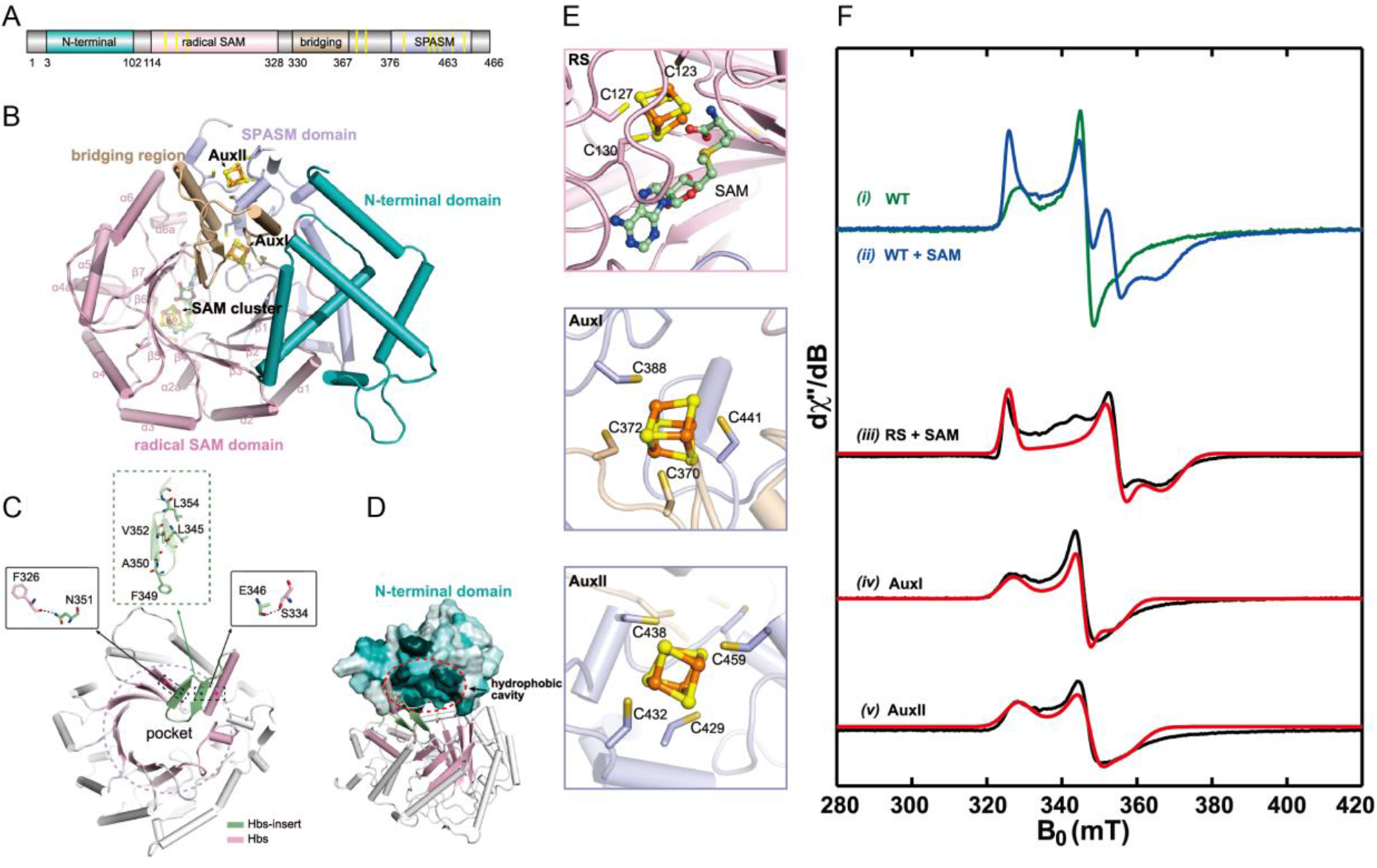
EPR spectroscopic characterization and AlphaFold2 predicated structure model of Hbs. (A) Four domains shown in Hbs. (B) Overall architecture of Hbs, showing an N-terminal domain comprised of six helices, a conserved RS domain, a bridging region with two helices and a β-harpin motif, as well as a C-terminal SPASM domain. (C) The interaction between the β-hairpin and the pocket involves two hydrogen bonds between F326 and N351, as well as S334 and E346. (D) Cavity map of Hbs active site reveals a hydrophobic pocket. (E) Three [4Fe-4S] clusters in the structure model of Hbs. (F) X-band CW EPR spectra of various Hbs samples (i) the wild-type (WT) Hbs reduced by DTH; (ii) subsequently adding SAM to DTH reduced Hbs; (iii) the mutant sample of Hbs (C438AC370AC372A) containing only radical SAM (RS) cluster reduced by DTH, and incubated with SAM; (iv) the mutant sample of Hbs (C438AC123AC127A) containing only AuxI cluster reduced by DTH; (v) the mutant sample of Hbs (C123AC127AC370AC372A) containing only AuxII cluster reduced by DTH. The black traces are experimental spectra, and the red traces are simulated spectra.

The radical SAM domain of Hbs encompasses three quarters of the triosephosphate isomerase (TIM) barrel fold and features a site-differentiated [4Fe-4S] cluster, which is a shared characteristic among all radical SAM enzymes (Fig. 2B). Hbs exhibits a substrate-binding pocket similar to Tes, but this pocket is significantly reduced in size due to the insertion of the β-hairpin within the bridging region, rendering it incapable of accommodating the GDGTs substrate adequately (Fig. S3). The interaction between the β-hairpin and the pocket primarily involves two hydrogen bonds between F326 and N351, as well as S334 and E346 (Fig. 2C). On the opposite side of β-hairpin, there are multiple hydrophobic amino acids. Therefore, it is hypothesized that during the GDGTs substrate binding, this hairpin flips outwards, providing sufficient space for substrate binding and stabilizing the substrate binding through its hydrophobic side (Fig. S3). Additionally, the hydrophobic cavity from the N-terminal domain of Hbs, which faces the substrate-binding pocket (Fig. 2D and Fig. S3) may also help to stabilize the substrate binding (Fig. 2D and Fig. S3). Combining both structural model analysis and FeS cluster identification by EPR spectroscopy, we proposed a molecular mechanism for the C(sp^3^)-C(sp^3^) linkage between two isoprenoid chains of GDGTs to form H-GDGTs (Fig. S3).

### The biological sources of H-GDGTs

To identify the biological sources of H-GDGTs, we performed a BLASTP search (e-value< 1e^-50^, identity>30%, sequence length>420 amino acids)of archaea genomes in the NCBI database for the Hbs protein homologs. We identified 649 Hbs homologs distributed widely in all three archaeal superphyla (Asgard, Tack, and DPANN) and the Euryarchaeota (Fig. 3A). However, it is intriguing to note that all Hbs homologs are exclusively present in obligate anaerobic archaeal genomes.

**Fig. 3.**
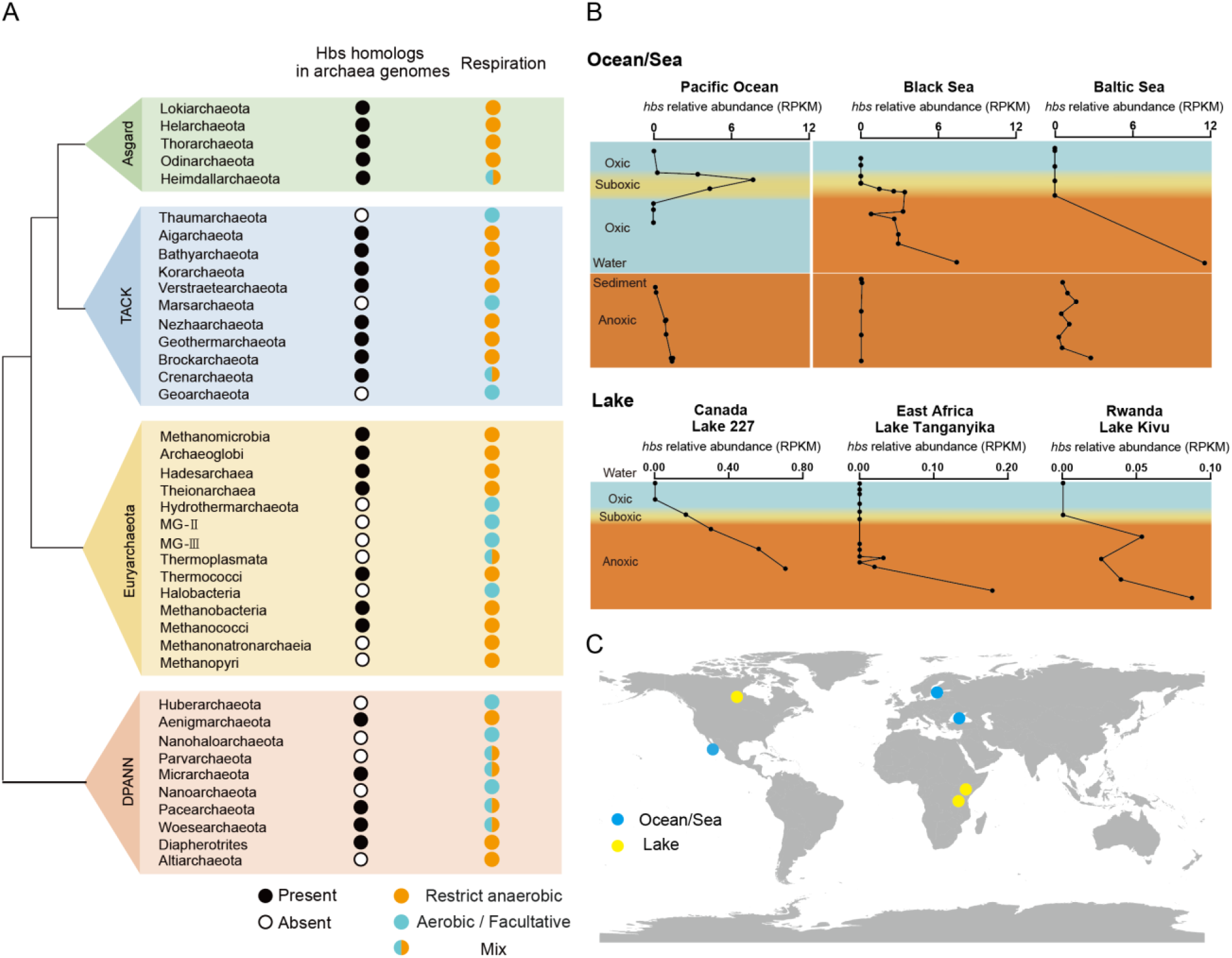
The distribution of Hbs homologs in archaea genomes and environmental metagenomes. (A) The BLASTp searches of Hbs protein homologs in the NCBI database and the respiration features of archaea are derived from published studies (*27-39*). Taxonomic groups that contain at least one species with Hbs homologs are designated by black circles. It’s important to note that the presence of black circles in a group does not imply that every species within the group possesses Hbs homologs. Conversely, taxonomic groups lacking any Hbs homologues are represented by white circles. For respiration features, the groups contain only obligate anaerobic archaea are marked in orange, aerobic and facultative archaea were marked in blue, and the groups contain anaerobic and aerobic/facultative archaea are marked in mix color of orange and blue. (B) The relative abundance of *hbs* gene homologs is present as RPKM values, and the reads of *hbs* were obtained by tBLASTn searches of metagenomes in SRA database on NCBI. The sample depths and oxygen concentration profiles are included in Data File. The color indicates oxygen levels (blue is oxic, yellow is suboxic, and orange is anoxic). (C) Geographical distribution of analyzed metagenome samples.

The lipid analysis of culturable archaea from previous studies was further applied to explore the potential link between H-GDGTs production (Hbs) and anaerobic respiration feature. Some obligate anaerobic archaea, including *Methanothermobacter thermautotrophicus, Palaeococcus helgesonii, Pyrococcus furiosus, Thermococcus acidaminovorans, Aciduliprofundum boonei*, and *Ignisphaera aggregans*, contain Hbs homologs and were reported to produce H-GDGTs (*10, 19-21*). In contrast, aerobic archaea, such as marine thaumarchaea, one of the most abundant archaea in marine water column, do not have Hbs homolog and thus, do not produce H-GDGTs (*22*). Moreover, even the facultative anaerobic archaea, such as *Metallosphaera sedula* and *S. acidocaldarius*, lack both Hbs homolog and H-GDGTs lipids (*23, 24*) as well. Thus, a preponderance of evidence exists to support that the biosynthesis of H-GDGTs is exclusive to obligate anaerobic archaea.

### *hbs* gene present in oxygen-deficient environments

The distinct biological source of H-GDGTs suggests that these lipids most likely link to oxygen-deficient ecosystems and environments. We next investigated the presence of *hbs* gene homologs in the metagenomes extracted from oxic or oxygen-deficient (defined as O_2_<3μM) (*25*) environments with tBLASTn searches (e-value<1e^-20^, identity>60%, sequence length>80 bp), which indicates the microbial communities have the genetic capacity to synthesize H-GDGTs. The relative abundance of *hbs* was calculated as RPKM (reads per kilobase of sequence per million reads) values.

Ocean metagenome sequences, in conjunction with oxygen level information, were obtained from the SRA (Sequence Read Archive) database on NCBI. These sequences were extracted from Pacific Ocean, Black Sea and Baltic Sea, encompassing various oceanic environments including the ocean surface, suboxic water (oxygen minimum zone, OMZ) and anoxic marine sediment/water. These metagenome sequences were subsequently analyzed for the presence of *hbs*. As expected, *hbs* was exclusively detected in the suboxic/anoxic (oxygen-deficient) environments, but absent in oxic waters (Fig. 3B). In general, water columns tend to exhibit a higher relative abundance (RPKM values) of *hbs* compared to the sediment samples, suggesting H-GDGTs were primarily synthesized within aquatic environments. However, *hbs* was not found in every oxygen deficient sample, such as certain samples from Baltic Sea suboxic water and Black Sea sediment (Fig. 3B). This implies fluctuations in H-GDGTs producers within oxygen-deficient environments could be influenced by other environmental factors.

In addition to the ocean, we observed a comparable distribution pattern of *hbs* in freshwater lakes (Fig. 3B). We specifically choose representative lakes with available metagenome data and oxygen level measurements for our analysis. These lakes included Lake 227 in Canada, Kivu Lake in Rwanda, and Lake Tanganyika in East Africa. Unlike the open ocean, the lower waters of most lakes are anoxic even euxinic due to limited water circulation (*26*). In these lake environments, *hbs* homologs were found exclusively in oxygen-deficient waters. The widespread presence of *hbs* homologs across diverse oxygen-deficient environments highlight H-GDGTs as a versatile tool for investigating oxygen deficiency in various environmental settings.

### H-GDGTs a new proxy for ocean deficiency in the present and past

One significant advantage of using lipid biomarkers for paleoenvironmental studies is their enduring preservation in the geological records. We first examined the relative abundance of H-GDGTs lipids in modern seas, alongside direct measurements of oxygen levels (*40-42*). The findings from the three sites in the Baltic Sea and Black Sea revealed that that H-GDGTs were much more prevalent in oxygen-deficient waters compared to oxic waters (Fig. 4A). These observations were consistent with the distribution of *hbs* genes in various environments. Taken together, it implies the H-GDGTs lipid profiles can reflect ocean oxygen depletion.

**Fig. 4.**
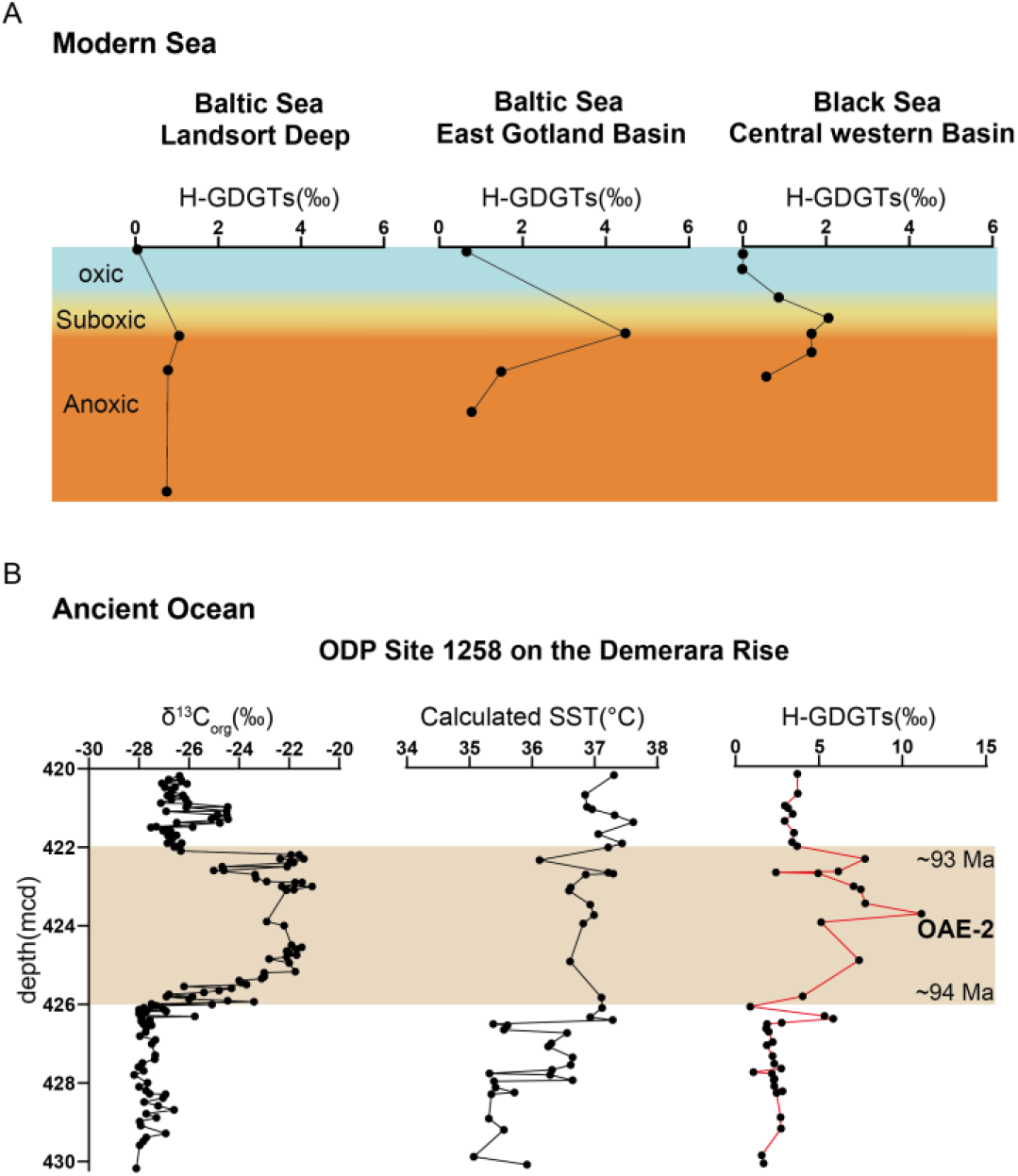
The distribution of H-GDGTs lipids in modern and ancient environments. (A) H-GDGTs were predominantly present in modern oxygen-deficient seawaters. The sample depths and oxygen concentration profiles are included in Data File. The color indicates oxygen levels (blue is oxic, yellow is suboxic, and orange is anoxic). (B) Relative amount of H-GDGTs was accumulated in sediments during the OAE-2 interval (ODP Site 1258, 422-426 mcd). The δ^13^Corg (%) and Calculated SST data are from Connock et al. (*45*).

The relative abundance of H-GDGTs over total GDGTs is calculated using equation 1:

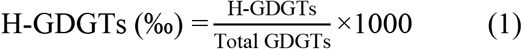

where total GDGTs include H-GDGTs, GDGT 0-8 and crenarchaeol.

To evaluate the viability of H-GDGTs as proxy for tracing historical ocean oxygen deficiency, we measured the H-GDGTs relative abundance in Cenomanian-Turonian sediments (∼94 Ma) from the Demerara Rise in the southern North Atlantic Ocean (ODP Site 1258) spanning Oceanic Anoxic Event 2 (OAE-2). The results show a remarkable increase in H-GDGTs (‰) during OAE-2 period (Fig. 4), aligning with the excursion of δ^13^Corg (%) which is established marker to characterize global OAEs (*43*). Furthermore, the H-GDGTs (‰) are slightly higher in the post-OAE-2 interval (420-422 mcd) compared to the pre-OAE-2 interval (426-430 mcd), a trend consistent with δ^13^Corg (%) records. More specifically, the H-GDGTs proxy provides a more detailed and explicit depiction of the ocean oxygen deficiency in the interval that bound the OAE-2, as the enrichment of H-GDGTs was sustained until the onset of OAE-2 (426.40-426.30 mcd) and is approximately contemporaneous with a thallium isotope excursion (*44, 45*), indicating the regional deoxygenation occurred before the globally recognized carbon cycle perturbation. Additionally, the variations of H-GDGTs (‰) were not correlated with the calculated sea surface temperature (SST) (Fig. 4B), suggesting the production of H-GDGTs was not notably influenced by changes in SST. These findings imply H-GDGTs have the potential to serve as a proxy for studying ocean anoxic events dating back hundreds of millions of years in Earth’s history.

## Discussion

The direct linkage of two sp^3^ carbons is a challenging biochemical reaction. Similar bond formation was reported in Tes (or MJ0619) (*5, 6*), and GrsA/B (*7*). All of these enzymes are radical SAM enzymes, which utilize radial SAM chemistry to accomplish two sequential C(sp^3^)-H activation on the inert sp^3^-hybridized carbon centers. Then the question is how these enzymes store the high-energy radical intermediates, such as the first substrate carbon radical generated via H-atom abstraction by the 5’-dAdo•. For Tes (or MJ0619), an intermediate with a bond between the substrate carbon and a sulfur of one auxiliary [4Fe-4S] was identified, providing evidence that the substrate radical is stabilized by coupling with the [4Fe-4S] cluster (*6*). However, for radical SAM enzymes GrsA and GrsB, which catalyzes the cyclization reaction of GDGT-0 at C7 and C3 position, respectively. Based on the sequence analysis, it most likely contains only one radical SAM [4Fe-4S] cluster. Therefore, GrsA and GrsB probably employ another way to stabilize the substrate radical intermediate, as there is no auxiliary cluster. In this work, we identified that Hbs is a SPASM-domain containing radical SAM enzymes, which harbors one radical SAM [4Fe-4S] cluster and two auxiliary [4Fe-4S] clusters. Therefore, it likely catalyzes C-C bond formation through stabilizing the substrate carbon radical via a C-S bond with the auxiliary [4Fe-4S] cluster, similar to Tes.

Why is H-GDGT exclusively produced by anaerobic archaea? The Hbs enzyme, which belongs to radical SAM superfamily, catalyzes the reaction under strongly reducing condition. This fact alone cannot explain why only anaerobic organisms produce H-GDGTs, as many other radical SAM enzymes are also extensively used by aerobic organisms. The reducing environment within the cell cytoplasm may be sufficient for radical SAM enzyme to function even under aerobic culture. Alternatively, Hbs shares homologous with AhbC and AhbD, which are responsible for the anaerobic biosynthesis of heme b in methanogens and sulfate-reducing bacteria, an alternative route to synthesis heme in contrast to the classical pathway found in eukaryotes and most bacteria (*46, 47*). The homology and evolutionary relationship of these three radical SAM enzymes suggest Hbs was possibly inherited from an anaerobic ancestor, providing a plausible explanation for the exclusive presence of H-GDGTs in anaerobic archaea.

H-GDGTs holds the promise of serving as a novel ocean anoxia biomarker, delivering a more comprehensive understanding of past ocean oxygen deficiency. The presence of H-GDGTs primarily depends on shift in its associated microbial communities in water column, which are highly sensitive and responsive to the rise of anoxic condition. Thus, unlike carbon isotope characterization, which reflects the large-scale perturbations of the global carbon cycle (*48*), the H-GDGTs proxy has the advantage to yield a more precise and regional representation of ocean oxygen deficiency supported by its well-established biological basis. However, the use of H-GDGTs proxy may be limited due to the low abundance of H-GDGTs in certain marine sediment samples. The relative abundance of H-GDGTs to total GDGTs ranges from 0.9 to 11‰ in ODP Site 1258 from Demerara Rise in this study, approximately 4‰ in ODP201-1229A-22H1, but they were not detected in ODP201-1229D-4H4 from Peru margin (*49*). Besides, the deep-sea hydrothermal vent systems have the potential to contribute to H-GDGTs production due to the thriving populations of thermophilic anaerobic archaea (*50-52*). Despite sharing anoxic conditions, hydrothermal vent regions differ significantly from regular oceanic anoxic water regions where oxygen depletion primarily results from the decomposition of organic matter. Hence, caution should be exercised when applying H-GDGTs records to interpret the cause of ocean oxygen deficiency in the presence of deep-sea hydrothermal vent influence.

In summary, our study has unveiled the biosynthetic gene, identified the biological sources, and elucidated the synthetic mechanism of H-GDGTs. These findings not only expand our understanding of lipid biosynthesis and chemistry, but also enable the development of H-GDGTs as a novel biomarker for indicating diverse oxygen-deficient environments, particularly for tracking the history of ocean deoxygenation.

## Acknowledgements

We thank Weiqi Yao and Lu Fan from Southern University of Science and Technology for constructive discussion, and Miao Huang from China University of Geosciences (Wuhan) and Fengfeng Zheng from Southern University of Science and Technology for the help in lipid analysis. Support for this study was provided by National Nature Science Foundation of China to Z.Z (No.32170041 and 92051112); Guangdong Innovative and Entrepreneurial Research Team Program to F.Z. (No. 2021ZT09Y104).

## Author contributions

Z.Z, L.T and F.Z. were responsible for the conceptualization and supervision of the project; Y.L., L.T., T.Y., F.Z., B.Z., H.C., H.R.A and N.Z.B. conducted genetic, biochemical and structural experiments; X.F., Y.L. and Z.Z. performed bioinformatic analysis; X.L., H.Y., J.K., G.T.C., A.K.W., L.D. and F.W. provided environmental samples and performed lipid analysis; Y.L., Z.Z., L.T. and F.Z. wrote the manuscript with input from all of the authors.

